# Integrated cross-sectoral surveillance of antimicrobial resistance genotypes and phenotypes across disparate reservoirs

**DOI:** 10.1101/2025.10.29.685460

**Authors:** Thomas D. Watts, Thanavit Jirapanjawat, Laura Perlaza-Jiménez, Francesco Ricci, Eve Tudor-Matthew, Eleonora Chiri, Sean K. Bay, Rhys Grinter, Rachael Lappan, Trevor Lithgow, Laura C. Woods, Chris Greening

**Affiliations:** Infection Program, Biomedicine Discovery Institute and Department of Microbiology, Monash University, Clayton, VIC, Australia; Centre to Impact AMR, Monash University, Clayton, VIC, Australia; Securing Antarctica’s Environmental Future, Monash University, Clayton, VIC, Australia

## Abstract

Antimicrobial-resistant (AMR) bacteria and genes are continually exchanged among humans, animals, and environmental reservoirs. Disparate and siloed surveillance methods present a major challenge for tracing and disrupting AMR transmission, with clinical monitoring focusing on detecting specific pathogens or specific genes of interest, and environmental surveys often relying on inferences drawn from indicator organisms. Here, we demonstrate that, following sample-specific pre-processing, common surveillance approaches can be applied consistently to profile AMR abundance, distribution, and phenotypes across diverse reservoirs, including soil, sediment, water, wastewater, and faecal samples from both urban and agricultural settings. Across all sample types, three core methods provided complementary insights: (i) quantitative PCR (qPCR) arrays to measure multiple AMR genes, (ii) gene- and genome-centric metagenomics for comprehensive resistome profiling, and (iii) culture-based genomics with susceptibility testing to link genotypes to phenotypes. We applied this approach to profile 1,032 metagenome-assembled genomes, 66 bacterial isolate genomes, and 78 and 6,442 AMR genes/reference sequences via qPCR and metagenomics, respectively. This integrated framework revealed a moderate prevalence but high diversity of resistance mechanisms in both pathogens and non-pathogenic bacteria with potentially transmissible genes, with wastewater especially enriched in AMR genes. We detected mismatches between genotype and phenotype predictions and a prevalence of intermediate resistance phenotypes, highlighting how many mechanisms of environmental resistance remain poorly understood. Overall, this study demonstrates that unified field-leading surveillance methods can be extended beyond clinical contexts into diverse environmental and animal samples, while highlighting that multiple methods are needed to capture the diverse AMR genotypes and phenotypes in these settings to enable comprehensive monitoring and adaptive solutions to restrict transmission.

## Background

Without intervention, AMR-associated infections are predicted to cause up to 10 million deaths annually by 2050, alongside a trillion-dollar economic burden on global healthcare systems (Murray et al. 2022; O’Neill 2016). AMR is quintessentially a One Health issue, with resistance determinants and resistant bacteria consistently exchanged between humans, animals, and the environment (Larsson and Flach 2022; Lappan et al. 2024). The concept of resistome describes all the resistance genes present in a given microbial community (D’Costa et al. 2006). Assessing the sources and impacts of environmental resistomes is difficult as the natural diversity of ARGs in the environment, making it hard to interpret what’s “normal” and what is indicative of a problem such as imminent spillover to another reservoir. Additionally, resistance may emerge due to selective pressures enriching antimicrobial resistance genes (ARGs) in resident microbiota, contamination with bacteria from human or animal sources that harbor existing resistance genes, or a combination of both (Bengtsson-Palme and Larsson 2015; Larsson and Flach 2022). For example, in many countries antibiotics used in human medicine are also extensively applied in livestock production (Granados-Chinchilla and Rodríguez 2017; Martin et al. 2015; Butaye et al. 2003), promoting AMR in animals with the potential for cross-reservoir transmission. Resistant bacteria are also released into the wider environment via wastewater discharge, septic leakage, stormwater runoff, animal faeces, and agricultural runoff (Rizzo et al. 2013; Karkman et al. 2018). Urban-adjacent water systems can act as important reservoirs and transmission routes, particularly in low-middle income settings (Bauza et al. 2020). Sub-lethal concentrations of antibiotics from agricultural, hospital or wastewater sources also act as selective pressures in these systems, fostering the persistence and diversification of resistant populations (Gullberg et al. 2011).

In these environments, AMR can occur through acquisition of mutations, or the acquisition of ARGs via horizontal gene transfer mediated by mobile genetic elements. Horizontal gene transfer has the potential to embed resistance traits in resident microbial communities, or spreading ARGs from environmental bacteria to clinically relevant human pathogens (Wyres and Holt 2018; Wang et al. 2024; Goh et al. 2024). Bacteria with broad niches, such as drug resistant *Klebsiella pneumoniae,* have been highlighted as conduits allowing environmental ARGs to enter clinical settings (Wyres and Holt 2018). Importantly, environmental and agricultural settings are not only a passive sink, but active incubators of resistance, where several clinically important ARGs including plasmid-derived *mcr-1* (colistin resistance), were first identified in animals before becoming widespread in the clinic (Liu et al. 2016). Identifying hotspots of ARG diversity and abundance is also critical for understanding the emergence of novel resistance genes (Bengtsson-Palme and Larsson 2015; Böhm et al. 2020).

Despite the importance of environmental reservoirs, AMR surveillance remains disproportionately pathogen-centric and biased toward clinical settings. The most recent Antimicrobial Usage and Resistance in Australia (AURA) report highlights that surveillance still focuses on clinical and aged-care facilities, focusing only on AMR pathogen detection (AURA 2023). In addition, environmental surveillance often relies on single indicator organisms to signal faecal contamination, representing a narrow approach that poorly reflects the diversity and abundance of pathogens or AMR in environmental niches (Harwood et al. 2005; Anderson et al. 2005; Ishii et al. 2006). Clinical AMR monitoring is tightly regulated and globally coordinated, by contrast, approaches for environmental surveillance vary, with a range of sample types necessitating tailored approaches, detection methods are diverse, there is limited agreement on which organisms and ARGs should be prioritised, and no global standardised frameworks exist (Bengtsson-Palme et al. 2023). Furthermore, many surveillance efforts use either genotypic or phenotypic methods in isolation. However, as we recently reviewed (Lappan et al. 2024), different techniques each have complementary strengths and limitations: targeted qPCR enables sensitive detection of known ARGs, but cannot identify novel variants; metagenomics provides broad gene discovery, but limited resolution of host context; and culture-based isolation and genomics links ARGs to phenotypic resistance, but captures only a fraction of microbial diversity in an environment. Additionally, linking ARG detection to clinically relevant phenotypes is also important. Gene carriage does not always translate to resistance, and without confirming phenotype, the clinical significance of environmental AMR can be uncertain. A unified, cross-sectorial surveillance framework that integrates complementary methods is therefore essential to move beyond documenting AMR, and towards building actionable systems to inform mitigation and intervention strategies across the One Health spectrum (Berendonk et al. 2015; Huijbers et al. 2019; Bengtsson-Palme et al. 2023; Lappan et al. 2024).

Here, we demonstrate the feasibility and utility of an integrated approach to investigate the distribution of AMR across diverse environmental reservoirs within the greater Melbourne metropolitan area. Using coordinated methodologies including qPCR, metagenomics, bacterial isolation, antibiotic susceptibility testing, and whole genome sequencing we generate a comprehensive view of ARG carriage and resistance phenotypes across soil, water, and faecal samples. This unified framework enables high-resolution identification of environmental AMR and helps inform future surveillance strategies that can be applied in a range of contexts.

## Results

### Antibiotic resistance genes and pathogenic genera are present within disparate environmental sources across Melbourne, Australia

We sought to understand the distribution of AMR pathogens in the environment surrounding Melbourne and to compare different surveillance approaches. Ten diverse sample types were selected, including agricultural runoff and sludge collected from wetland water, urban park soil and water, animal scats, wastewater, urban beach sediment, river water, and a dairy farm (**Table S1**, **Figure S1**). Samples were collected and pre-processed differently depending on sample type, but all subsequent procedures including DNA extraction and downstream analysis were conducted using the same pipeline. To assess the distribution of ARGs within the different environments, we performed qPCR arrays to detect 78 genes associated with AMR or virulence (**Figure 1**). Wastewater is a rich environmental sink where microbes from diverse sources converge, and the highest diversity of ARGs were associated with this sample type (average 55 ARGs). Genes from almost all antimicrobial classes tested by the qPCR were represented in wastewater samples, including determinants of beta-lactam resistance (*bla_OXA_*, *bla_SHV_*, *bla_KPC_*, *bla_IMP_*, *bla_VIM_*, *bla_NDM_*, *bla_CTX-M_*), vancomycin resistance (*vanA* and *vanB*), erythromycin resistance (*ermA*, *ermB*, *ermC*), tetracycline resistance (*tetA*, *tetB*), and fluoroquinolone resistance (*qnrA*, *qnrB*, *qnrD*, *qnrS*) (**Figure 1**).

**Figure 1.**
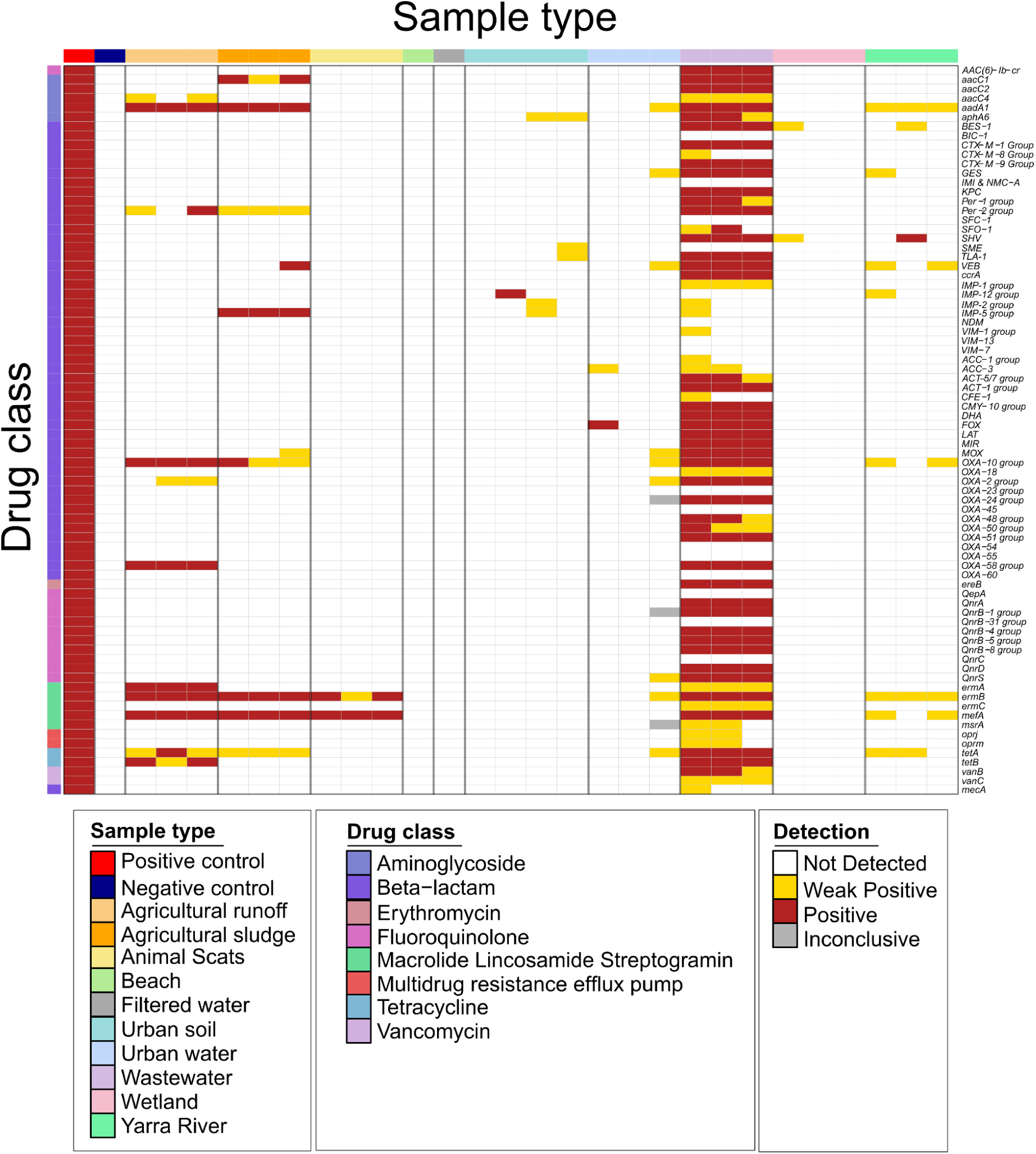
ARG detection by qPCR array. DNA extracts were applied to the QIAGEN Microbial DNA qPCR array for Antibiotic Resistance Genes, which encompasses a range of ARGs. A “+/−” result indicates weak detection (qPCR threshold value within 3-6 cycles of the no-template control).

ARGs were broadly distributed in other samples, although there were far fewer ARGs detected per sample (average 3.6) compared to wastewater (average 55.6). Agricultural samples showed ARGs associated with tetracycline resistance, macrolide resistance, beta-lactam resistance, macrolide resistance, and aminoglycoside resistance. Animal scats collected from an urban park were also positive for the determinants of macrolide resistance. Beach samples were negative for all ARGs tested, whereas urban park soil and water contained determinants of aminoglycoside resistance and beta-lactam resistance. Wetland and river samples were positive for aminoglycoside, macrolide, beta-lactams, and tetracycline resistance determinants.

### Metagenomic datasets mirror AMR distribution highlighted by qPCR arrays and link ARGs to specific metagenome assembled genomes

Although qPCR rapidly reports the diversity of known ARGs that are present in the environment, it cannot identify ARGs other than those included in the array. qPCR also has limited capacity to link ARGs to specific hosts which is important for understanding emerging resistance and reservoirs. We therefore conducted deep metagenomic sequencing (NextSeq100) on each sample to complement qPCR surveillance, generating an average of ∼240 million reads per sample (120 million paired reads). Identical methods for library preparation, sequencing, and downstream analysis were used for all samples. Quality controlled reads from each metagenome dataset were used to query the CARD database (implemented via the RGI tool) to determine the breadth of ARGs present (**Figure 2**). This catch-all approach yielded a more comprehensive picture of the total resistome of these samples. Unlike qPCR results, all samples appeared to have a rich diversity of ARG classes, with wastewater samples displaying the highest number of drug classes, and the highest average abundance of ARGs (∼32-33 classes, average abundance of ARGs in the community 1.2% ± 0.2% SD) (**Table S4**). The increase in the number of ARG classes identified for each sample likely reflects the enhanced breadth of detection afforded by using a more comprehensive database (6,642 genes). Especially when compared with the targeted approach of qPCR (78 genes), where the specific primers used in the qPCR array preclude the capture of the full diversity of sequences within each ARG group. Resistance to aminoglycoside, disinfectants and antiseptics, tetracycline, fluoroquinolone and glycopeptide were both widespread and highly abundant in all sample types. Beta-lactam resistance, including carbapenem resistance, were also present within all samples.

**Figure 2.**
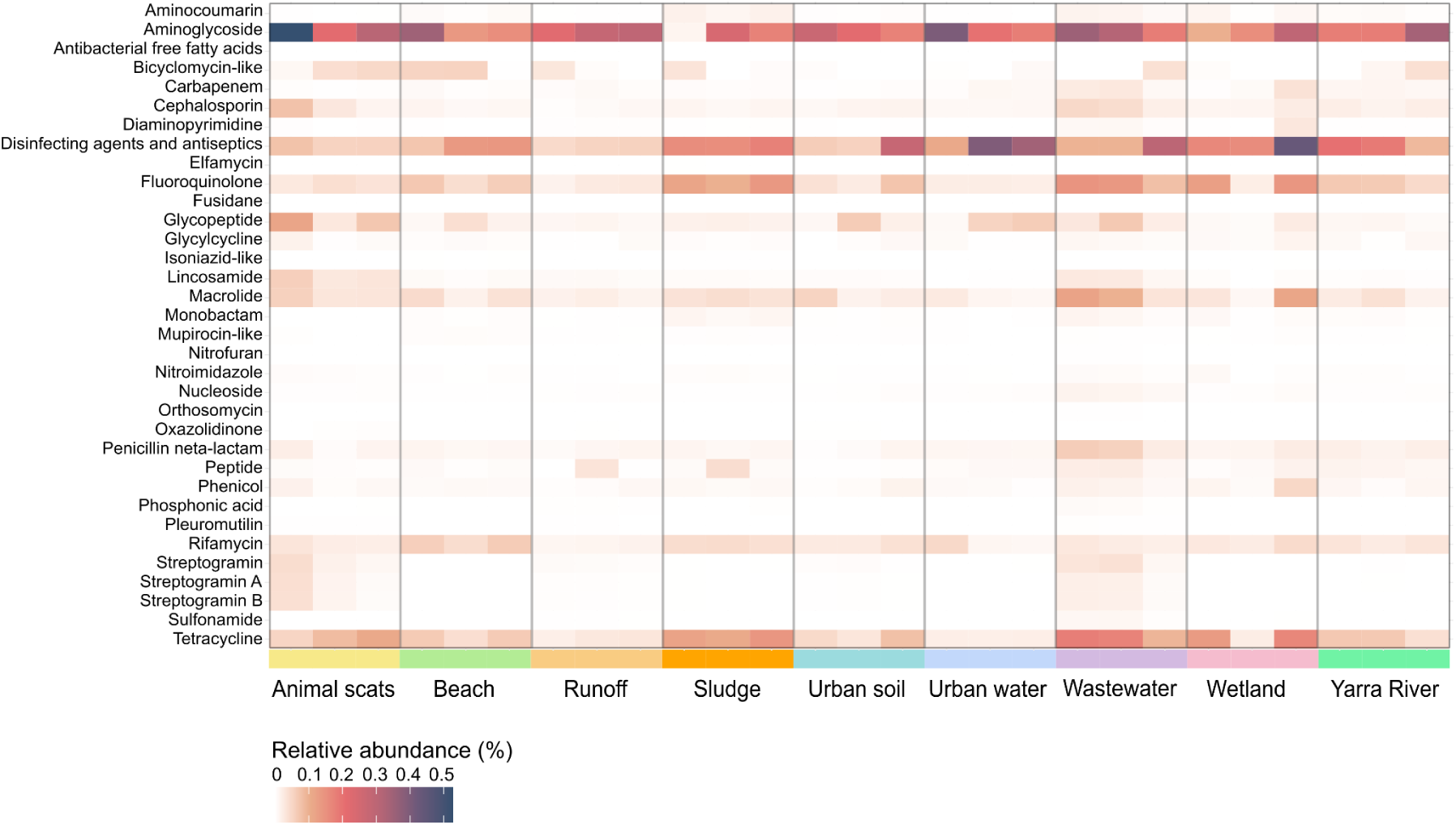
Heatmap depicting ARG abundance in metagenome reads with the CARD resistance gene identifier (RGI). Metagenome short reads were subsampled to 20 million read pairs for each sample type (n=3) and RGI was used to determine ARG carriage. Detected ARGs were collapsed to drug class and relative abundance (%) was calculated for each sample.

To better link ARGs/AMR to specific community members and identify potential emerging AMR pathogens or reservoirs, metagenomes within each environment were co-assembled and binned to yield 1,032 species level (0.95 ANI) metagenome assembled genomes (MAGs). Mapping reads from each sample to cognate MAGs showed an average coverage of 18.79% (**Table S2**).

Taxonomic classification of MAGs showed that they span 27 bacterial phyla encompassing Pseudomonadota/Proteobacteria (24.7%), Bacteroidota (19.9%), Bacilliota/Firmicutes (19.5%), and Actinomycetota (17.3%) (**Table S3**). The majority of MAGs appeared to represent autochthonous organisms commonly associated with their environment, although putative opportunistic human pathogens were identified in a subset of samples (**Table S3**), including *Campylobacter upsaliensis* (animal scats) (Goossens *et al*., 1990), *Escherichia coli* (beach sand, constructed wetland, river), and *Aeromonas sobria* (Spadaro *et al*., 2014) (urban water). Genomes of several other potential pathogens were also reconstructed from wastewater, namely *Streptococcus suis* (Hughes et al. 2009), *Sutterella wadsworthensis* (Kirk et al. 2021), *Bilophila wadsworthia* (Gan et al. 2025; Baron et al. 1992), *Laribacter hongkongensis* (Abot et al. 2020; Engsbro et al. 2018), and *Aliarcobacter butzleri* (Prouzet-Mauléon et al. 2006).

MAGs were screened for ARGs using comprehensive databases (CARD), which were subsequently collapsed into AMR class and used to adorn a maximum-likelihood phylogenetic tree (**Figure 3a**). ARGs were present in each environment and were not confined to a single phylum (27 phyla). Genes associated with resistance to glycopeptides (*vanG, vanH, vanO, vanT, vanW, vanY*) (n=619, 60%), tetracyclines (*tetA, tetB, adeF, tetQ, tetZ, otrA, poxtA, tet(36)*) (n=231, 22.4%), fluoroquinolones (*adeF, marA, soxS, soxR, sdrM, rsmA, acrA, acrB, acrE acrF, acrS, emrA, emrB, emrR, mdtH, patB*) (n=201, 19.5%), and disinfecting agents/antiseptics (*qacG*, *qacJ*, *qacL*) (n=79, 7.7%) were most prevalent within MAGs, including numerous non-pathogenic bacteria (**Figure 3a, Table S3, Table S5**).

**Figure 3.**
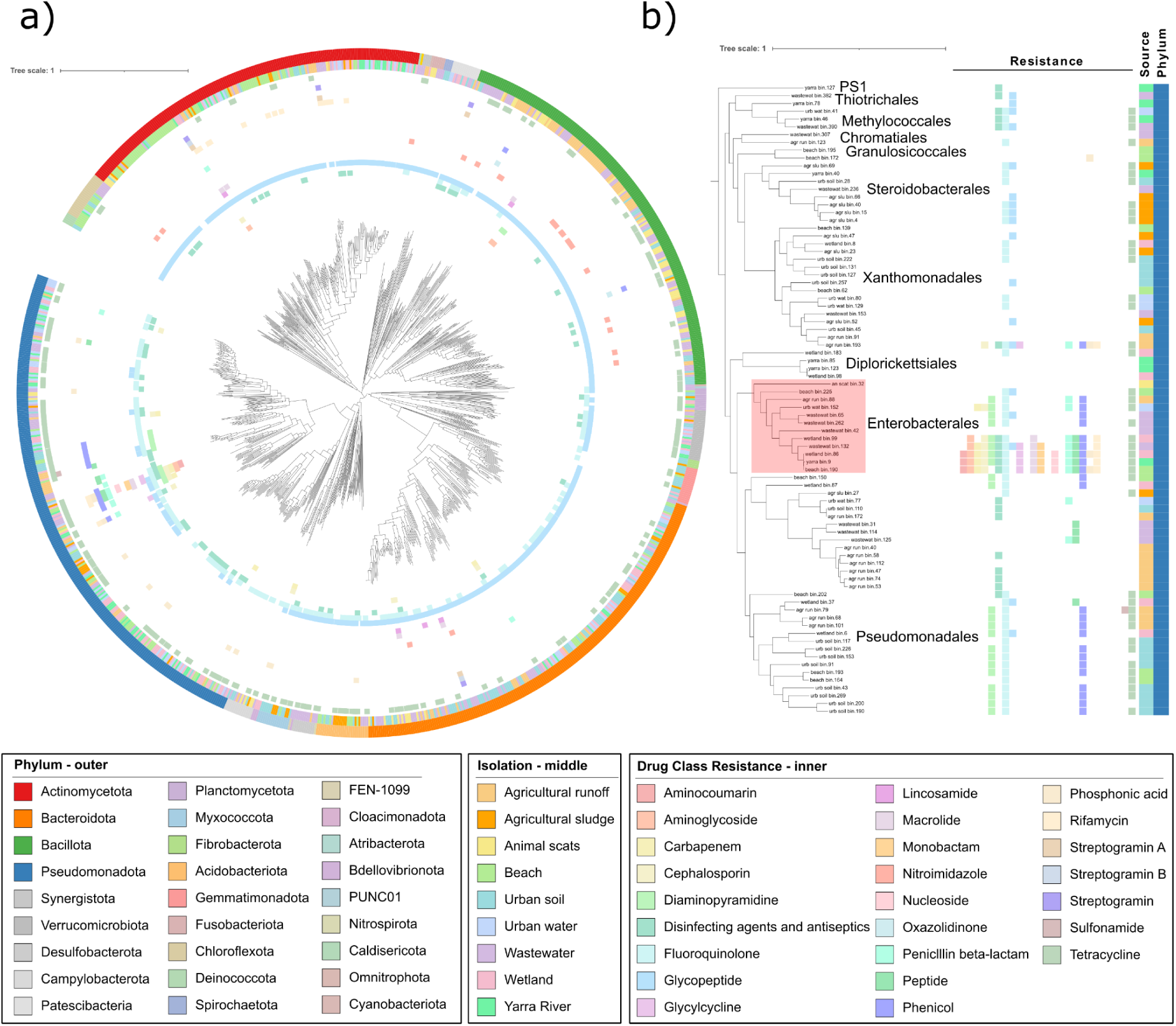
ARGs show a widespread distribution within MAGs but are prevalent within Pseudomonadota. a) Maximum-likelihood phylogenetic tree of 1,032 environmental MAGs reconstructed from samples taken across Melbourne, Australia. Outermost ring indicates phyla, middle ring indicates isolation source and innermost rings indicate drug resistance determined via comparison to the CARD database with RGI. b) A zoomed inset showing Pseudomonadota MAGs with high ARG carriage.

Resistance to carbapenem and cephalosporin was more limited (n=9, 0.9% and n=12, 1.2% respectively), but still detected among both autochthonous bacteria (*Mangrovibacter spp., Flavobacterium spp., Qipengyuania spp., Pantoea spp*., *Phocaeicola vulgatus*, *Bacteroides uniformis*, *Luteimonas spp.*, *Rhodoferax* spp.) and putative pathogens (*E. coli, L. hongkongenesis, A. sobria*). Pseudomonadota MAGs had the most AMR diversity, with some genomes harbouring genes conferring resistance to up to 17 different antibiotics (**Figure 3b**). Notably three *E. coli* MAGs from beach, wetland and river sources possessed putative resistance to 16-17 antibiotic classes, including carbapenems. Other bacteria with high resistance capacity included a *Pantoea spp.* (12 drug classes), *Mangrovibacter spp.* (11 drug classes) and the putative pathogen *A. sobria* (7 drug classes).

### Isolates with intermediate resistance phenotype are present in almost all sample sites

To support our culture-independent investigation, we selected bacterial isolates from each source for antibiotic susceptibility testing and genomics analysis, with the aim of determining whether AMR genotype is concordant with phenotype. Following collection and dilution, samples were plated onto selective media (MacConkey agar, Bile-Esculin agar and Sheep Blood agar) without antibiotic selection to isolate putative pathogens and other environmental organisms, before subsequent identification with 16S rRNA gene sequencing. Following culturing, quality control, and presumptive identification through standardised methods, a total of 66 isolates were identified for downstream analysis (**Table S6**). All samples contained putative pathogens (**Figure 4a**), suggesting they are widely distributed across the target environments. Isolates predominantly belonged to Pseudomonadota, with some also from Actinobacteria and Firmicutes (**Figure 4b**). Few isolates, including *Escherichia coli* and *Acinetobacter pseudolwoffii,* were represented in the MAGs (**Table S3, Table S6**). This finding reflects that MAG construction only captured an average of 18.8% of the community and is biased towards abundant bacteria, whereas culture-based approaches select primarily for fast growing or prolific organisms such as pathogens, including those in very low abundance. Indeed, mapping metagenome reads to isolate genomes showed only 0.55% coverage (**Table S2**).

**Figure 4.**
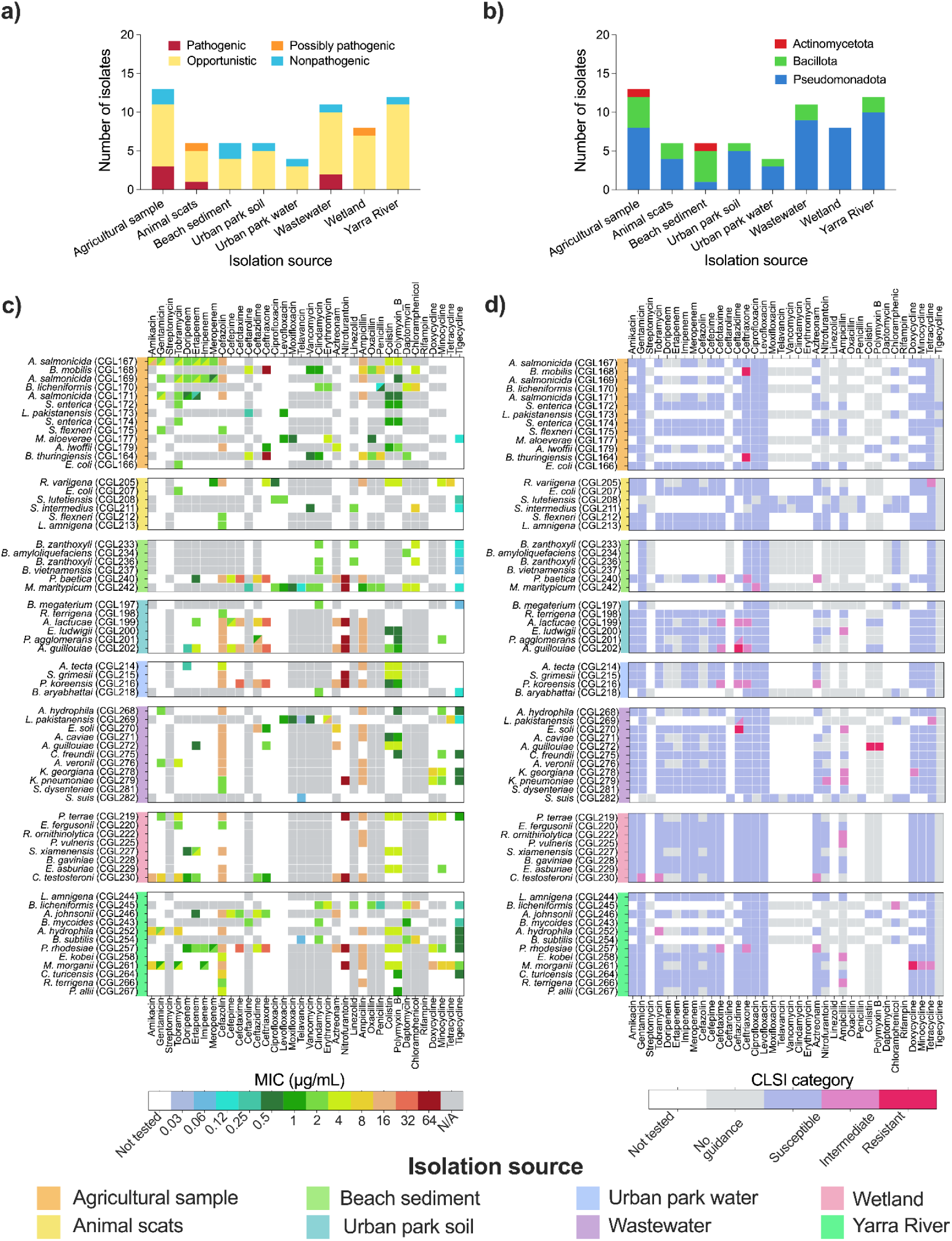
Resistance profiles of isolates by minimum inhibitory concentration (MIC) and as interpreted using CLSI thresholds (CLSI, 2018). Isolates of sufficient quality (see Methods) are categorised by **a)** pathogenicity and **b)** phylum. **c**) MIC for each isolate grouped by sampling site **d**) Resistance of each isolate as determined by the CLSI guidelines. Cases in which resistance is ambiguous are depicted as cells split along the diagonal, with upper left and lower right indicating the lowest and highest resistance level observed, respectively.

To determine the resistance profile of each environmental isolate, Minimum Inhibitory Concentrations (MIC) for a range of antibiotics were determined, where antibiotics were tested based on isolate Gram status (**Figure 4c**). Nine isolates (13.6%) were resistant to antibiotics based on the Clinical and Laboratory Standards Institute (CLSI) guidelines using the most appropriate threshold based on taxonomy of each organism, although for many taxa, guidance was not available for many antibiotics and therefore resistance could not be determined (**Figure 4d**). For example, CLSI does not currently include specific guidelines for tigecycline, underscoring the need for better understanding of antibiotic resistance in environmental contexts. All samples indicated at least one isolate showing an intermediate resistance (also known clinically as ‘susceptible, increased exposure’) phenotype, suggesting that these environments could potentially act as reservoirs for emerging resistance. The number of isolates with intermediate to high resistance according to the CLSI 2018 differed between samples, ranging from 15.4% (agricultural site) to 67.7% (urban park soil). Numbers are affected by sampling bias, but also CLSI guidance availability for isolated organisms. The isolate with resistance to the highest number of antibiotics was *Morganella morganii* (CGL261), which was resistant to seven different antibiotic classes, including the third-generation antibiotic imipenem (**Figure 5c**). Two other opportunistic pathogens, *Citrobacter freundii* (CGL275) (Anderson et al. 2018) and *Proteus tarrae* (CGL219) (Zheng et al. 2022; Li et al. 2022), were each resistant to four antibiotics. While these species are of concern due to their potential pathogenicity to humans, in most cases, resistance was conferred by chromosomal genes.

**Figure 5.**
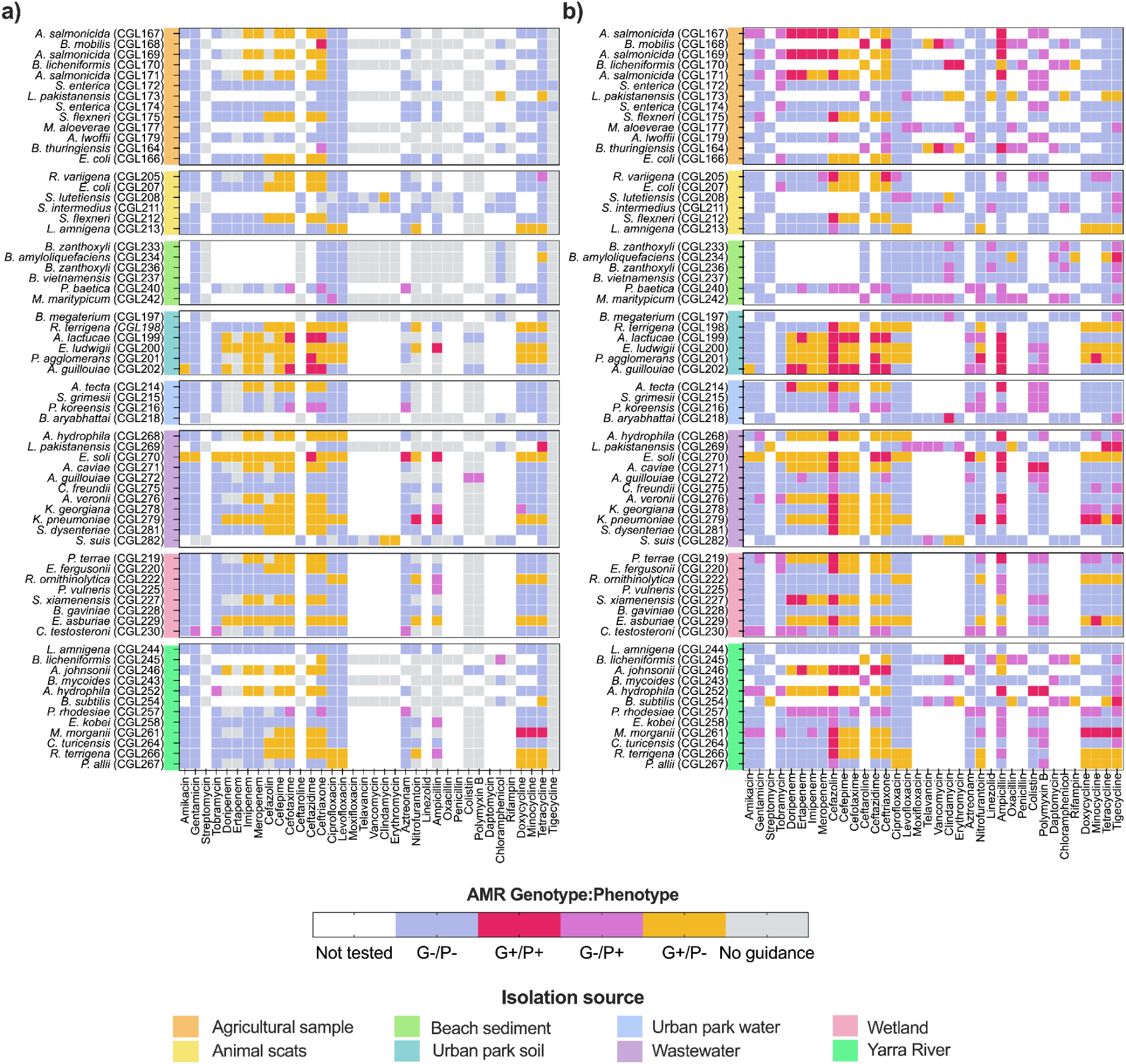
Genotype-to-phenotype contingency plot for antibiotic resistance in environmental isolates. Sampling sites are shown on the left axis of the figure. In **a)**, CLSI (2018) thresholds are used to interpret resistance. Resistance here includes intermediate resistance and full resistance as defined by the CLSI (2018). In **b)**, resistance is defined as any MIC above 0 μg/mL to include low-level resistance.

### Isolate AMR genotype and phenotype are often discordant

To examine if the AMR phenotypes recorded for each isolate were in accordance with ARG carriage, whole-genome sequencing (WGS) was conducted for each isolate and the CLSI and MIC data was overlaid. Considering the CLSI guidelines for resistance, nine isolates showed evidence of agreement between their resistance phenotype and the presence of a corresponding ARG. All isolates showed evidence of susceptible phenotypes when ARGs were not present as expected (**Figure 5a**). However, many isolates also showed susceptible phenotypes, despite carrying an ARG that is predicted to confer resistance to the corresponding antibiotic class. Additionally, several isolates showed a resistant phenotype with no evidence of cognate ARG carriage, suggesting that these isolates either encode uncharacterised resistance genes or have intrinsic antibiotic-tolerance mediated by a non-genetic factor. As the CLSI guidelines are not suitable for all bacteria isolates, we also examined resistance using MIC data, where we considered an MIC over 0.1 μg/mL as ‘resistant’, in order to include low-level resistance phenotypes (**Figure 5b**). Taking MIC into consideration, many of the isolate-antibiotic combinations that previously were either not represented by the CLSI guidelines or showed a susceptible/ARG present phenotype were somewhat resistant. Additionally, the MIC data also indicated more strains were had low levels of resistance despite not encoding an ARG; the assumption that an MIC of > 0 μg/mL indicates resistance likely increases reporting of false positives, although these isolates could also be evidence of emerging AMR or antibiotic tolerance. ARGs were identified for each isolate (**Table S7**), where the most common class of ARG detected in these isolates were those conferring resistance to cephalosporins (34.6%) and beta-lactams (25.6%). However, most of the isolates carrying genes conferring this antimicrobial resistance were in fact susceptible to these antibiotics when assayed.

A critical issue to address was the extent to which the ARGs are plasmid-associated ARGs, given the ready transmission of phenotypes via plasmids. The data showed that most ARGs were chromosomally encoded, suggesting limited capacity for dissemination. Although these chromosomal genes could be associated with other mobile genetic elements such as transposons (**Figure S2**). All resistance genes conferring intrinsic resistance were located on the chromosome, while genes associated with acquired resistance were encoded both on plasmids and the chromosome. Only 12.1% of isolates carried ARGs on plasmids, notably *Aeromonas caviae*, *Aeromonas hydrophila*, *Lysinibacillus pakistanesis*, *Enterobacter soli*, *Kluybera georgiana*, *Acinetobacter johnsonii*, and *Rahnella variigena* isolates, and several plasmids were potentially mobilizable (have *mob* or relaxase genes) indicating they have potential to be transferred by co-resident conjugative plasmids (**Figure S2**).

## Discussion

This study demonstrates that approaches typically applied in clinical AMR surveillance can be successfully extended to environmental and agricultural contexts. By deploying a coordinated workflow encompassing qPCR, metagenomics, and culture-based genomics across diverse reservoirs, we demonstrate that unified methodologies can profile AMR across the One Health spectrum. This approach contrasts with conventional environmental surveillance, which has often relied on indicator organisms or single-method approaches that underestimate both the diversity and distribution of resistance determinants (Berendonk et al. 2015; Lappan et al. 2024). Clinically aligned methods can be adapted to capture resistance dynamics across environmental compartments at a resolution comparable to hospital-based monitoring, providing a practical model for future One Health surveillance.

We recommend applying multiple methodological pillars for surveillance, as their complementary strengths are required to paint a more complete picture of the environmental resistome. qPCR arrays delivered rapid and sensitive quantification of clinically relevant ARGs, but still provided a narrow perspective (even with 78 genes quantified) and minimal contextual information (e.g. which microbes carry ARGs). Metagenomics broadened this view, revealing a wider spectrum of ARGs, including many harboured by non-pathogenic taxa that would have been overlooked by targeted methods. Many of these ARGs overlap with clinically important resistance genes, mirroring other studies that show environmental and agricultural-derived ARGs detected by metagenomics and metatranscriptomics overlap extensively with clinically important antibiotic resistance genes (Shi et al. 2025). Co-assembly and binning of our data produced over 1,000 species-level MAGs, capturing abundant autochthonous taxa, while also identifying putative opportunistic pathogens. Despite this depth, MAGs provided only a partial perspective on low-abundance pathogens and mobile genetic elements, highlighting the continued importance of more targeted approaches, including culture-based phenotyping. The traditional approach of culture-based isolation remains best practice for functional validation and allows genotype to phenotype links to be tested, though this is biased to only a subset of organisms and excludes the environmental majority that are recalcitrant to culturing, which was further highlighted by the low mapping rate observed when aligning metagenome reads to isolate genomes. Combining these approaches also highlighted discrepancies between predicted and observed resistance, including intermediate phenotypes. In some cases, ARGs were detected without corresponding resistance, likely reflecting differences in gene expression, reading frame, copy number, or genomic background (Wagner et al. 2023). Conversely, resistance without identifiable genetic determinants was also observed, potentially pointing to the contribution of intrinsic mechanisms to antibiotic tolerance, or the existence of novel or uncharacterised resistance genes. Examples of this include non-cabapenemase dependent carbapenem-resistance phenotypes in *Klebsiella* that depend on changes in the outer membrane permeability to provide intrinsic resistance to carbepenems (Rosas et al. 2023). Together, these findings underscore that carriage of ARGs does not always equate to phenotypic resistance, and that phenotype can emerge independently of known genotypes.

The CLSI guidelines (CLSI, 2018) provide a clinically relevant accounting of resistance for applicable species, which remain indispensable for patient management and clinical surveillance. However, their application to environmental and non-clinical taxa is less straightforward. The behaviour of a resistance gene can vary considerably depending on its mode of transmission, copy number, and the genomic background of the host, meaning that CLSI interpretive breakpoints may not be useful predictors of emerging resistance. At the same time, persistence of ARGs without conferring phenotypic resistance is possible and may obscure the expected link between genotype and phenotype (Martínez et al. 2015). This is why well-controlled MIC testing remains essential for accurately characterising phenotypic resistance in environmental isolates. Nevertheless, persistence of “silent” ARGs can still pave the way for the emergence of resistance (Wagner et al. 2023). Together, these factors highlight both the value and the limitations of applying clinical interpretive frameworks to environmental resistance data and underscores the need for parallel ecological and functional perspectives.

Currently there are no equivalent frameworks for interpreting environmental resistomes. Developing an effective framework for understanding AMR risk requires a nuanced approach that accounts for factors such as the host, ARG mobility, local selective pressures and the co-occurrence of primary or opportunistic pathogens that could receive ARGs from environmental hosts. Without these contextual factors, it remains difficult to assess whether a given resistome profile represents a benign feature of the environment or an emerging risk. Several studies have taken steps to incorporate some of these components into their surveillance strategies by measuring selective agents including antibiotic residues and selective factors such as heavy metals or biocides, integrating these data with paired metagenomic analysis (Lundström et al. 2016; Bengtsson-Palme et al. 2019). However, most studies still capture only part of the picture. Our combined approach helps to expose these weaknesses, demonstrating how much interpretative power is lost when surveillance depends upon a single method or perspective. This contributes to the broader goal of building an interpretive framework for surveillance of environmental AMR that is tailored to complex sample types and microbial communities.

Overall, we observed a relatively low prevalence but high diversity of resistance determinants. This finding likely reflects both the relatively high-income setting of Melbourne (established waste management protocols to reduce bacterial load etc). Wastewater was a critical reservoir, containing an array of clinically important determinants, reinforcing the value of wastewater epidemiology (Hendriksen et al. 2019). Beyond known pathogens, ARGs were frequently identified in endemic species such as those of *Pantoea* and *Mangrovibacter*, many of which carried genes spanning multiple antibiotic classes. This result reinforces previous observations that environmental bacteria can act as reservoirs of resistance, even when not pathogenic, highlighting their potential role in horizontal gene transfer to clinically relevant species (Forsberg et al. 2012). These results provide additional high-resolution evidence of ARG carriage in non-pathogenic hosts across both urban and agricultural environments, demonstrating that even in a relatively low-prevalence setting, diverse resistance mechanisms occur outside clinical settings. Future work could now extend this framework across broader geographic and socioeconomic contexts, and through longitudinal monitoring, to better resolve how resistomes evolve under different selective pressures.

## Methods

### Collection of samples

Fourteen samples were collected (Supplementary Table 2) in sterile 50-ml centrifuge tubes and processed within 48 hours by resuspending 1 g in 9 ml of sterile peptone saline diluent (1 g/L peptone, 8.5 g/l NaCl, pH 7.0), and plating 10 µl of the supernatant onto three different selective agar plates: Sheep Blood agar (SBA), Bile esculin agar (BEA), and MacConkey agar (MAC) plates (Media Preparation Unit, Department of Microbiology & Immunology, The University of Melbourne). Samples were further diluted before plating as shown in Supplementary Table 2. Plates were incubated aerobically at 37°C for 24 hours. Several colonies with different morphologies and biochemical activity indicated by selective plates were chosen and re-streaked onto a fresh selective agar plate, and incubated at 37°C for 24 – 48 hours to further isolate the strain. A single colony from each selective agar plate was re-streaked onto Luria-Bertani agar plate (10 g/L tryptone, 5 g/L yeast extract, 5 g/L NaCl and 15 g/L agar) and incubated at 37°C for 24 – 48 hours. For long term storage, bacterial stocks were prepared by culturing isolates in 10 ml LB broth (10 g/L tryptone, 5 g/L yeast extract and 5 g/L NaCl) at 37°C with shaking at 200 rpm for 24 – 48 hours and stored in 25% v/v sterile glycerol at −80°C.

### Genomic DNA extraction

gDNA was extracted using a modified CTAB method. Briefly, 2-4 ml of cell culture were pelleted by centrifugation at 11,000 x *g* and supernatant was discarded, before 30 μL of lysozyme solution (50 mg/mL in Tris-HCL pH 8), followed by vortexing an incubation at 37°C for 1 hour. Following incubation, 70 μL of 10% SDS and 10 μL of 20 mg/mL proteinase K were added and incubated for 15 minutes at 65°C. Next, 100 μL of 5 M NaCl and 100 μL of CTAB solution (10% CTAB in 0.7 M NaCl) were added, vortexed and incubated for 10 minutes at 65°C. Next, 750 μL of chloroform was added, followed by gentle mixing and centrifugation at 11, 000 x *g* for 15 minutes. The aqueous phase was transferred to a fresh tube and precipitated using 0.6 volumes of isopropanol and incubated at −20°C for 30 minutes. Precipitate was collected via centrifugation at 13, 000 x g for 10 minutes before washing with 1 mL of ice-cold 75% ethanol, and final centrifugation at 13,000 xg for 5 minutes prior to air drying for 30 minutes. The DNA pellet was resuspended in 50 μL of UltraPure water and left at 4°C to ensure dissolution.

### Bacterial isolate identification

Bacterial isolates were identified based on colony morphology and biochemical reactions on the selective agar plates as well as Sanger sequencing of the 16S rRNA gene. Single colonies on the LB agar plates were picked for colony PCR by resuspending the cells in 100 µl of UltraPure DNase/RNase-Free Distilled Water (Thermo Fisher Scientific) and boiled at 100°C for 10 minutes. The tubes were cooled on ice for 10 minutes before centrifugation at 16000 ×g for 10 minutes. The 16S rRNA gene was amplified from the supernatant using universal 16S rRNA gene primers (27F (5’-AGAGTTTGATCCTGGCTCAG-3’) and 1492R (5’-GGTTACCTTGTTACGACTT-3’), with an expected amplicon length of ∼1400 bp. Individual reactions contain 1 × Taq MasterMix (New England Biolabs), 400 µM of each primer, and 2 µl of the DNA template mixed to a final volume of 50 µl with UltraPure DNase/RNase-Free Distilled Water. Cycling conditions for PCR amplification were as follows: 95°C for 2 min followed by 35 cycles of 95°C for 30 sec, 56°C for 30 sec, 68°C for 1 min, with the final cycle ending at 68°C for 5 min before cooling down to 4°C. The PCR products were loaded on 1% w/v agarose gel at 90V for 35 min. The gel was visualised with a UV transilluminator and the band of the expected size (∼1400 bp) was cut and extracted using the ISOLATE II PCR and Gel Kit (Bioline) before sending for Sanger sequencing at Micromon Genomics (Monash University). If DNA was not amplifiable, genomic DNA was extracted as described above and used as a template for 16S rRNA gene PCR.The sequences were compared against the Standard Nucleotide BLAST database (NCBI) using a percent identity threshold 97%.

### Antimicrobial susceptibility test

Sixty-six isolates were selected for antimicrobial susceptibility testing using Sensititre plates (Thermo Fisher). Gram-negative strains were evaluated using Sensititre Gram Negative GN6F and Sensititre Gram Negative GNX2F plates. Gram-positive strains were analysed on Sensititre Gram Positive GPALL3F plates (Sensititre Plate Guide Booklet). The cultures were streaked from glycerol stocks onto LB agar plates and incubated overnight at 37°C. A few colonies were selected from each plate and resuspended in phosphate-buffered saline solution (13.69 mM NaCl, 2.68 mM KCl, 10.14 mM Na_2_HPO_4_, 1.76 mM KH_2_PO_4_, pH 7.4) and adjusted turbidity equivalent to 0.5 McFarland standard (0.24 mM BaCl_2_• 2H_2_O in 1% v/v H_2_SO_4_) or OD_600_ ≈ 0.15 – 0.18 and transferred 1 – 30 µl (Table 2) to Mueller-Hinton broth (Oxoid, Thermo Fisher). Following adjustment, 50 µl of the culture was loaded into each well of the Sensititre plate, sealed with perforated sealing labels and incubated at 37°C for 18 – 24 hours. Interpretation of Sensititre results was performed in accordance with the CLSI & Antimicrobial Susceptibility Testing (AST) M100 guidelines.

### Whole genome sequencing of isolates

Genomic DNA from each of the 66 isolates were extracted as described above and sent for whole genome sequencing (the MHTP Medical Genomics Facility, Hudson Institute). Libraries were prepared using the Celero EZ DNA-Seq library prep kit. Paired-end short read (150 bp) sequencing was performed with the Illumina NextSeq2000 platform. Quality control of the short-read libraries was evaluated using FastQC v0.11.7 (Andrews 2015) and adapters were trimmed from the reads using FastX ToolKit v0.0.13 (https://github.com/agordon/fastx_toolkit). Genomes were assembled with Unicycler v0.4.7 (Wick et al. 2017), and CheckM v1.1.6 (Parks et al. 2015) and QUAST v5.0.2 (Mikheenko et al. 2018) were used to evaluate the quality of the assemblies. GTDB-Tk v2.32 was used to assign assemblies to the GTDB (r214) taxonomy (Chaumeil et al. 2022; Parks et al. 2022). Isolates were removed from the analysis if their GTDB-assigned taxonomy differed from 16S rRNA gene PCR identification above the rank of genus, or if they could not be classified by GTDB-Tk likely due to contamination. A final set of 66 isolates were maintained for downstream analysis and had congruent taxonomy assignments at the rank of genus (**Table S6**).

### Plasmid and antibiotic resistance gene identification

Plasmids were predicted using PlasClass 06.2020 (Pellow et al. 2020) with a cut-off of 0.7 to classify contigs from draft assemblies as well as the function MOB-recon from MOB-Suite (Robertson and Nash 2018). Contigs for which PlasClass and MOB-Recon results were incongruent were manually examined and aligned with NCBI BLAST to the non-redundant nucleotide database to filter out contigs likely to be chromosomal. Prodigal v2.6.3 (Hyatt et al. 2010) implemented in Assembly2gene (https://github.com/LPerlaza/Assembly2Gene) was used to identify ARGs. A). Then the predicted genes were aligned with NCBI BLAST against an in house curated non-redundant ARGs database containing 6,657 genes, combining AMRfinderPlus (Feldgarden et al. 2021), the Comprehensive Antibiotic Resistance Database (CARD; (Alcock et al. 2020), ResFinder (Bortolaia et al. 2020), Beta-Lactamase DataBase (BLDB; (Naas et al. 2017), and membrane components such as porins and efflux pumps. Assembly2gene generates gene alignments and predicted peptides using R packages “msa”, “reshape2”, “Biostrings” and “seqinr”. Heatmaps and plots were generated using R packages “ggplot2” and PRISM (v9.0.0; Apple MAC, GraphPad Software, San Diego, California USA, www.graphpad.com). Putative ARG resistance mechanisms as provided by the CARD were compared with the MIC data to link isolate genotypes to their resistance profiles.

### Metagenomics

DNA was extracted from samples (**Table S8**) using the QIAGEN PowerSoil kit, as per manufacturer’s instructions with an elution volume of 30 μl. Samples were sequenced at the Australian Centre for Ecogenomics (ACE), using an Illumina NextSeq5000, to generate 2×150bp paired end reads (average of ∼238M reads per sample). Reads were assessed for quality and trimmed using fastp (Chen 2023), which included the removal of adapters and low-quality sequences, as well as filtering by read length (≥50 bp) and average quality score (≥30), with automatic adapter detection enabled. To enhance genome recovery, particularly of low-abundance taxa, reads from within each environment were co-assembled using MEGAHIT v1.2.9 using default parameters (Li et al. 2015). Assemblies were filtered to retain contigs ≥1,000 bp. Genome binning was independently performed with four algorithms: Vamb v4.1.3 (Nissen et al. 2021), MetaBAT v2.12.1 (Kang et al. 2015), CONCOCT v1.1.0 (Alneberg et al. 2014), and SemiBin2 (Pan et al. 2023). Bins were refined with the bin refinement module of MetaWRAP (Uritskiy et al. 2018). Bins were dereplicated with dRep v3.4.244 (Olm et al. 2017) at a 95% average nucleotide identity (ANI) threshold corresponding to species level, incorporating CheckM2 metrics. Completeness and contamination were assessed with CheckM2 (Chklovski et al. 2023). In total, we recovered 1,032 bacterial metagenome-assembled genomes (MAGs) meeting the MIMAG criteria for medium-(≥50% completeness; <10% contamination) and high-quality (≥90% completeness; <5% contamination). MAGs were taxonomically classified using GTDB-Tk v2.3.2 against the Genome Taxonomy Database (GTDB) release R214 (Chaumeil et al. 2022). A maximum-likelihood phylogenetic tree of bacterial MAGs was built using GTDB-tk v2.3.2 with hmmalign 3.4 model then visualised using iTOL v6 (Letunic and Bork 2024) and edited in InkScape. MAGs for each metagenomic sample were screened for ARGs using RGI. In addition, metagenome short reads were subsampled to 20 million read pairs using seqtk (v1.5) (https://github.com/lh3/seqtk) and also screened for ARGs using RGI, where relative abundance was calculated based on the number of reads mapped to ARG class divided by total number of reads (20 million).

