## Supplemental Figures S1 and S2 for "Integrated cross-sectoral surveillance of antimicrobial resistance genotypes and phenotypes across disparate reservoirs"

**Supplemental data:**

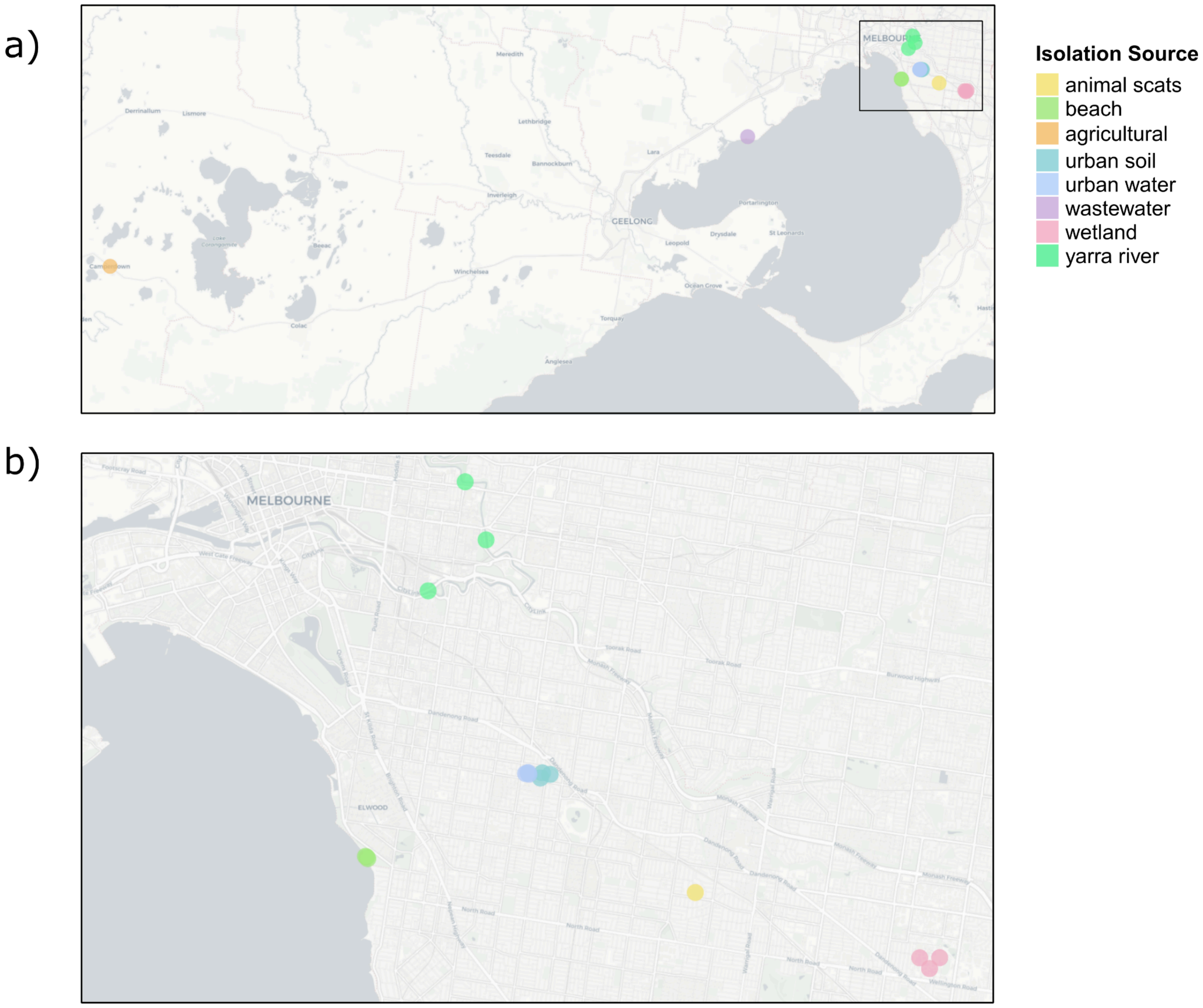


**Figure S1. Map of sample sites across Victoria, Australia.** a) wider sampling sites including agricultural run off and sludge collection at WestVic Dairy, Camperdown and wastewater at Western Treatment Plant. b) Zoomed view of eastern Melbourne sample sites.

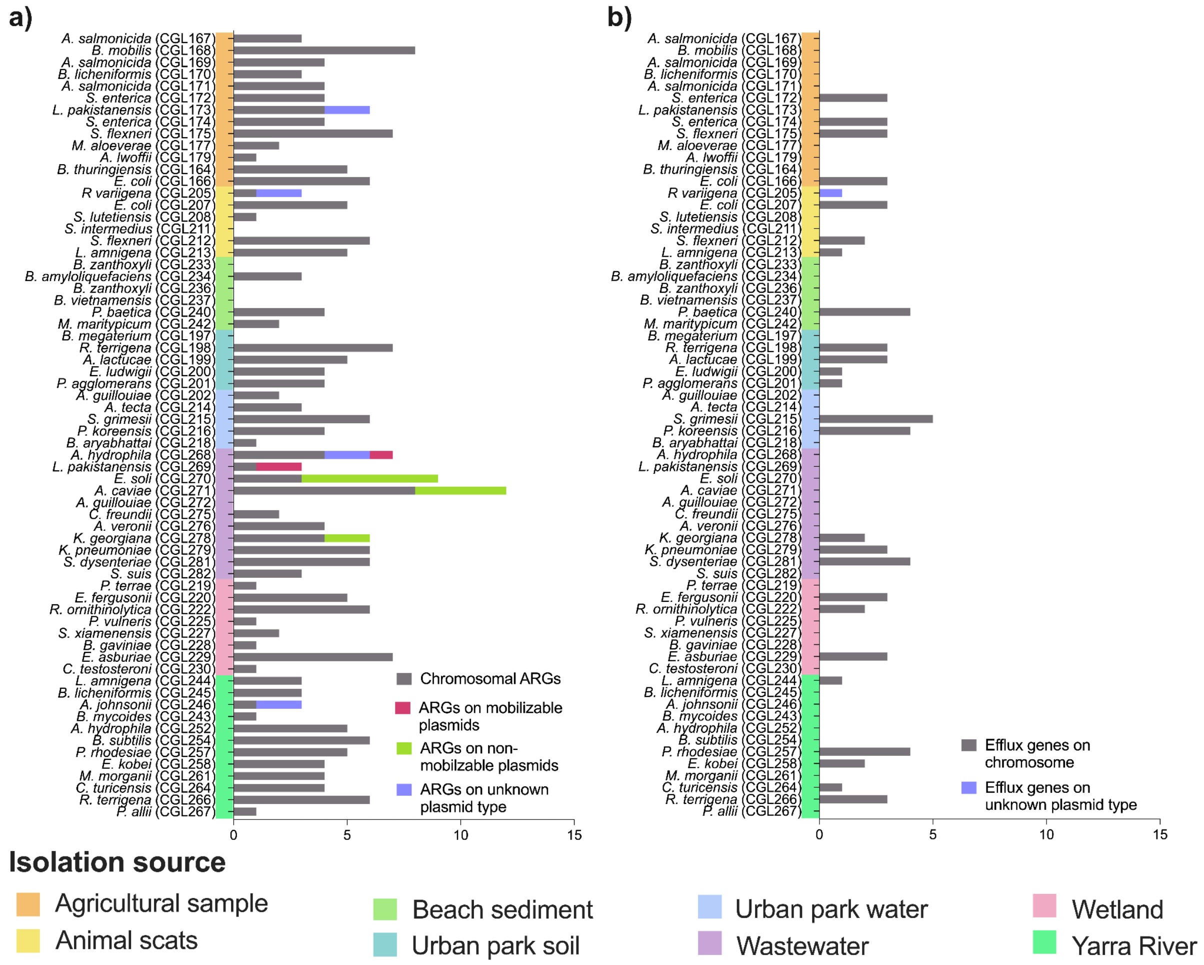


**Figure S2. Plasmid-borne genes among environmental isolates.** Non-efflux categorised antibiotic resistance genes (ARGs) identified with a percent identity of at least 70% are shown in **a)**, while efflux genes are shown in **b)**. The graph depicts the total number of ARGs present in each isolate separated into chromosomally encoded, or plasmid associated (mobilisable, non-mobilizable, and unknown).
